# Interdependency of respiratory metabolism and phenazine-associated physiology in *Pseudomonas aeruginosa* PA14

**DOI:** 10.1101/760454

**Authors:** Jeanyoung Jo, Alexa Price-Whelan, William Cole Cornell, Lars E.P. Dietrich

**Author notes:** Address correspondence to Lars E.P. Dietrich,.

## Abstract

Extracellular electron transfer (EET), the reduction of compounds that shuttle electrons to distal oxidants, can support bacterial survival when preferred oxidants are not directly accessible. EET has been shown to contribute to virulence in some pathogenic organisms and is required for current generation in mediator-based fuel cells. In several species, components of the electron transport chain (ETC) have been implicated in electron shuttle reduction, raising the question of how shuttling-based metabolism is integrated with primary routes of metabolic electron flow. The clinically relevant bacterium *Pseudomonas aeruginosa* can utilize carbon sources (i.e., electron donors) covering a broad range of reducing potentials and possesses a branched ETC that can be modulated to optimize respiratory efficiency. It also produces electron shuttles called phenazines that facilitate intracellular redox balancing, increasing the complexity of its metabolic potential. In this study, we investigated the reciprocal influence of respiratory metabolism and phenazine-associated physiology in *Pseudomonas aeruginosa* PA14. We found that phenazine production affects respiratory activity and terminal oxidase gene expression, and that carbon source identity influences the mechanisms enabling phenazine reduction. Furthermore, we found that growth in biofilms, a condition for which phenazine metabolism is critical to normal development and redox balancing, dramatically affects the composition of the *P. aeruginosa* phenazine pool. Together, these findings can aid interpretation of *P. aeruginosa* behavior during host infection and provide inroads to understanding the crosstalk between primary metabolism and shuttling-based physiology in the diverse bacteria that carry out EET.

**IMPORTANCE:** *Pseudomonas aeruginosa* is a major cause of healthcare-associated infections and long-term lung infections in people with cystic fibrosis. It can use diverse organic compounds as electron donors and possesses multiple enzymes that can transfer electrons from central metabolism to O_2_. These pathways support a balanced intracellular redox state and the production of cellular energy. Under hypoxic conditions, *P. aeruginosa* can reduce phenazines, secondary metabolites that also promote redox homeostasis and that contribute to virulence. We asked how these primary and secondary routes of electron flow influence each other. We found that phenazines affect respiratory function, that the roles of respiratory enzymes in phenazine reduction are highly condition-dependent, and that the complement of phenazines produced is strongly affected by growth in assemblages called biofilms. These results provide a more nuanced understanding of *P. aeruginosa* redox metabolism and may inform strategies for treating persistent infections caused by this bacterium.

---

Members of the domain Bacteria exhibit a vast variety of pathways and substrates that can be used to generate energy (1). This metabolic diversity allows bacteria to persist in conditions that do not support eukaryotes, which typically produce energy by redox transformations of oxygen and water (2). An additional advantage for many bacterial species is metabolic versatility, the ability to use multiple substrates and/or pathways for energy generation. Metabolic versatility provides resilience in heterogeneous or changing conditions and is an important feature of bacteria that form multicellular assemblages called biofilms.

Biofilms are clusters of microbes that adhere to each other by producing complex, polysaccharide-based matrices. In these structures, the combined effects of diffusion and metabolism generate resource gradients and unique conditions that differ from those found in historically popular, well-mixed liquid cultures. Importantly, biofilm-specific conditions can induce distinct physiologies that contribute to stress and antibiotic resistance, which in turn have important implications for virulence (3, 4). We seek to understand how redox metabolisms are integrated to support multicellular growth in the bacterium *Pseudomonas aeruginosa*, a common cause of chronic, biofilm-based infections.

*P. aeruginosa* uses organic compounds as carbon and energy sources. It can generate ATP by fermenting these compounds or by coupling their oxidation to generation of a proton motive force that powers ATP synthase (i.e., respiration). Results from our work, using a colony morphology model to study *P. aeruginosa* biofilm metabolism and development, support the model that electron flow to two major oxidants supports energy generation and redox homeostasis in biofilms: (1) O_2_ and (2) phenazines, redox-active metabolites that can shuttle electrons to oxidants available outside the cell and at a distance (**Figure 1**) (5, 6). Genetic analyses and microelectrode profiling studies indicate that *P. aeruginosa*’s ability to reduce O_2_ and phenazines depends on the composition of its electron transport (i.e., respiratory) chain (ETC) (7).

**Figure 1.**
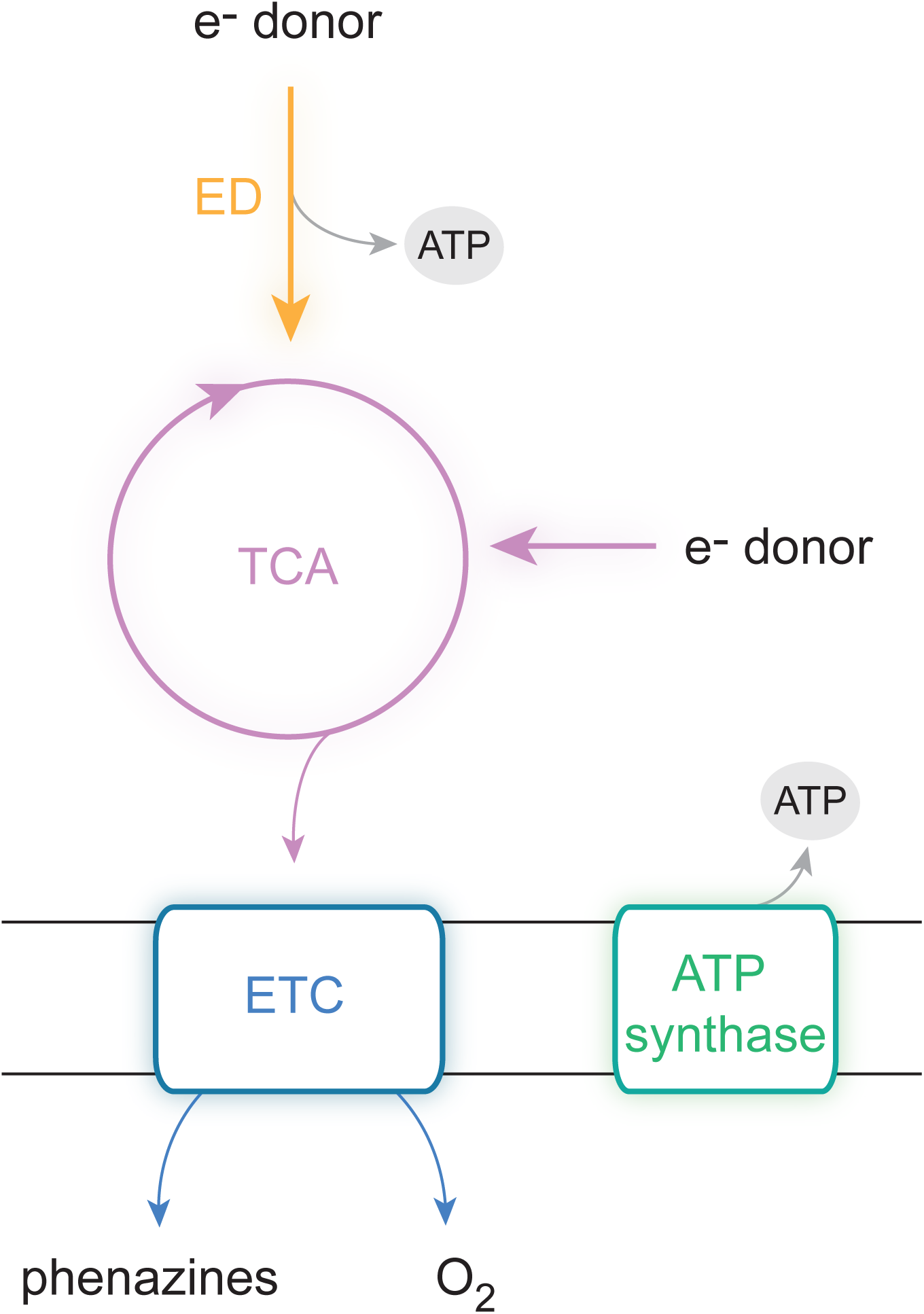
Routes of electron flow in *Pseudomonas aeruginosa* metabolism. Simplified representation of major redox pathways operating in *P. aeruginosa*. Orange, Entner-Doudoroff (ED) pathway; pink, the tricarboxylic acid (TCA) cycle; blue, aerobic respiration mediated by the electron transport chain (ETC). Electron transfer through the ETC is coupled to the generation of a proton motive force that powers the ATP synthase (green). ATP is also generated by the ED pathway. Electrons can originate from diverse carbon sources that can enter the ED pathway or TCA cycle.

In contrast to the mitochondrial ETC, which has one route for electrons to be delivered to O_2_, bacterial respiratory chains often have multiple routes (8). Electrons fed into *P. aeruginosa*’s ETC are ultimately transferred to one of five canonical terminal oxidases, the enzymes that catalyze the final electron transfer step to reduce O_2_ to water (9, 10). Each of these terminal oxidases possesses unique characteristics, including different expression patterns and affinities for O_2_ (11–14). Our picture of *P. aeruginosa*’s ETC is further complicated by the fact that its “Cco” complexes, which belong to the *cbb*_3_ family of terminal oxidases, can contain subunits encoded by multiple, redundant operons present at distinct sites in the genome (15). These heterocomplexes (i.e., isoforms) have specific roles under different conditions (15) and contribute differentially to biofilm physiology and redox state (7). In addition, work from our group has implicated specific Cco terminal oxidase isoforms in phenazine utilization in biofilms grown on a complex medium (7)

Though metabolism is often conceptualized as a modular process in which the oxidation of individual carbon sources can be paired with the respiration of electron acceptors via common, central pathways, this is an oversimplification. Different carbon sources (i) offer varying degrees of reducing power, (ii) are routed through specific pathways that will differ in their yields of direct electron donors to the ETC, and (iii) can directly or indirectly act as regulatory cues affecting the expression or activity of ETC components (16, 17). Furthermore, changes in electron flow through the ETC can affect the expression of terminal oxidase genes (18). Here, we examined the physiological impacts of phenazine production and ETC composition by surveying their relevance for respiratory activity during the use of diverse carbon sources in *P. aeruginosa*. We tested whether carbon source identity affects the ability to grow with specific respiratory chain components and the ability to produce and reduce specific phenazines during *P. aeruginosa* biofilm growth. Our results indicate that the identity of the provided carbon source has profound effects on PA14 growth and phenazine production and utilization. Furthermore, our data show that phenazines have differential effects on colony biofilm morphogenesis on distinct carbon sources and suggest that phenazines themselves can influence the expression of terminal oxidases within a biofilm.

## Results and Discussion

### *Phenazine production and ETC composition influence tetrazolium dye reduction in* P. aeruginosa

To assess how phenazine production affects *P. aeruginosa* PA14 respiration, we measured respiratory activity using commercially available “phenotype microarray” plates, which contain a different carbon source in each well. The manufacturer-provided growth medium for these plates contains a tetrazolium dye that is reduced by the electron transport chain (ETC) and thereby undergoes an irreversible color change (19, 20). First, we used phenotype microarray plates to compare tetrazolium dye reduction by wild-type (i.e., phenazine-producing) PA14 and the phenazine-null mutant Δ*phz*. We found that on four carbon sources--succinate, D,L-malate, acetate, and α-ketoglutarate (αKG)--phenazine production inhibited tetrazolium dye reduction (**Figure 2A**). On all other carbon sources respiratory activity was similar between WT and Δ*phz* (**Figure 3B**). During incubation in defined, MOPS-buffered media containing each of the 95 “phenotype microarray” carbon sources, 32 supported growth (**Supplementary File 1**), and WT and Δ*phz* grew similarly on all of these carbon sources (**Figure S1**). These results suggest that--during growth on succinate, D,L-malate, acetate, and αKG--phenazines divert electrons from the respiratory chain, which results in decreased tetrazolium dye reduction. All four of these carbon sources are intermediates in or branchpoints of the tricarboxylic acid (TCA) cycle, a major source of reducing power for respiration. αKG in particular is notable for its role as a precursor for glutamine and phenazine synthesis (21, 22).

**Figure 2.**
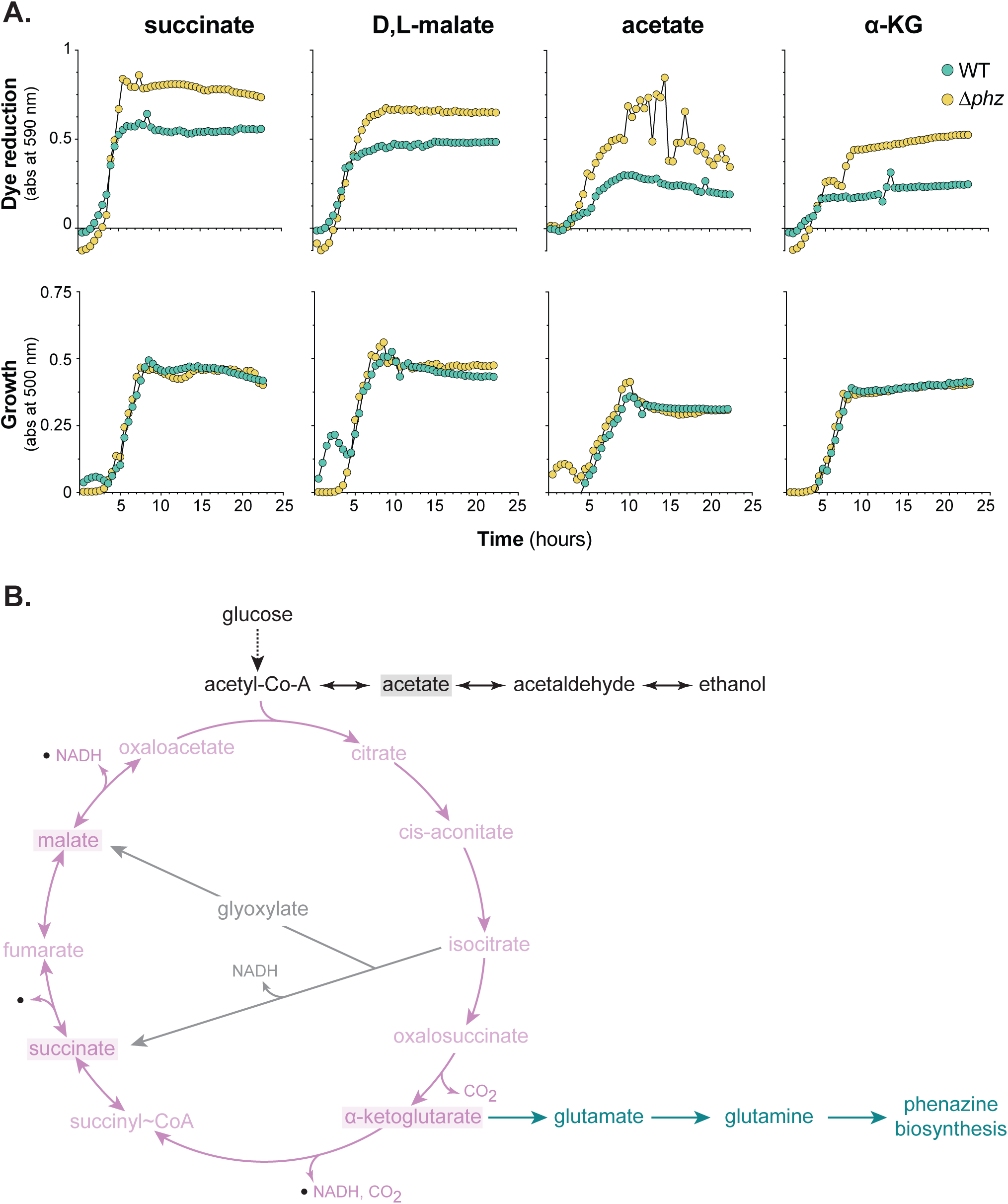
Phenazines influence tetrazolium dye reduction. **(A)** Dye reduction (top) and growth (bottom) of PA14 WT and the phenazine-null mutant (Δ*phz*) on succinate, D,L-malate, acetate, and αKG. Curves show data from one replicate, which is representative of data from three biological replicates; refer to **Figure S2** for full data set. **(B)** Schematic of the TCA cycle (pink) with the glyoxylate shunt (gray) and glutamine and phenazine biosynthesis (green) branchpoints indicated. Carbon sources identified in panel **2A** are shaded in boxes.

**Figure 3.**
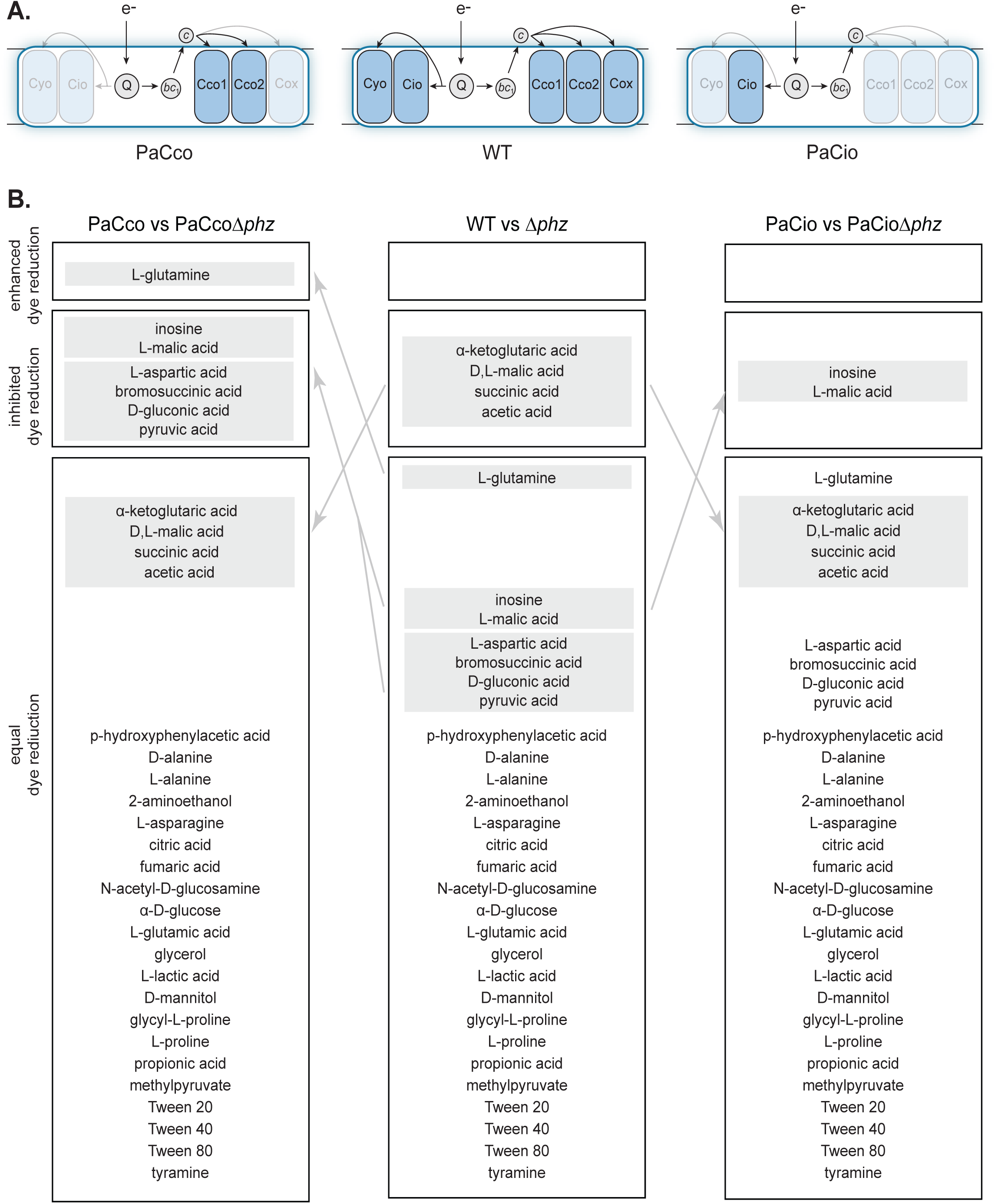
ETC composition influences tetrazolium dye reduction. **(A)** Depiction of the terminal oxidase complexes present in WT (center) and the terminal oxidase mutants PaCco (left) and PaCio (right). Electrons enter the ETC via primary dehydrogenases (not shown) and are transferred to the quinone pool (“Q”). Reduced Q can act as a substrate for the quinol oxidases (Cyo and Cio) or the cytochrome *bc*_1_ complex (“*bc*_1_”). A cytochrome *c* protein mediates electron transfer between *bc*_1_ and the terminal oxidases Cco1, Cco2, and Cox. **(B)** The 32 carbon sources that support PA14 growth, grouped based on tetrazolium dye reduction patterns of each phenazine-producing and phenazine-null strain. Groups were designated based on data from three experiments. Carbon sources highlighted in colored boxes indicate those on which dye reduction patterns changed relative to the strains with the full complement of terminal oxidases (i.e., WT and Δ*phz*; the center column). Carbon source descriptors are as provided by the manufacturer.

In a previous study, we found that electron transfer to phenazines depended on the composition of the ETC during growth in biofilms (7). We therefore suspected that the effects of phenazines on respiratory activity would also depend on the composition of the ETC. We generated mutants that contained only the two Cco’s (“PaCco”) or Cio (“PaCio”) (**Figure 3A**), because prior work has suggested that these terminal oxidases make the greatest contributions to aerobic and microaerobic growth in *P. aeruginosa* under laboratory conditions (10, 12). The Cco terminal oxidases accept electrons from a *c*-type cytochrome donor (downstream of the cytochrome *bc*_1_ complex and quinone pool), while the Cio terminal oxidase accepts electrons directly from the quinone pool (**Figure 3A**). Terminal oxidase gene mutations were created in wild-type and Δ*phz* backgrounds. We used phenotype microarray plates to measure respiratory activity, and also characterized the growth in a MOPS-buffered medium, of our terminal oxidase-mutant strains during incubation with each of the “phenotype microarray” carbon sources. While strains with all terminal oxidases (WT and Δ*phz*) and those with just the Cco’s (PaCco and PaCcoΔ*phz*) exhibited similar growth kinetics, the growth of PaCio strains was abrogated (**Supplementary File 1**), in line with previous findings that mutants lacking functional Cco terminal oxidases have growth defects (12, 14).

To test whether ETC composition influences the effects of phenazines on respiratory activity, we conducted phenotype microarray experiments for the PaCco and PaCio strains in phenazine-producing and phenazine-null backgrounds. We compared each phenazine-null mutant to its parent strain and categorized each carbon source depending on whether they enhanced tetrazolium dye reduction, inhibited dye reduction, or had no effect (**Figure 3B**). Because we encountered high variability in dye reduction between runs, we performed each experiment in biological triplicate and assigned a carbon source to a particular category if two out of the three trials produced similar dye reduction trends (**Supplementary File 2)**. During growth on most carbon sources, phenazine production did not affect respiratory activity, regardless of ETC composition (**Figure 3B**). However, we were intrigued to find that the phenazine-dependent inhibition of dye reduction--seen during growth on αKG, D,L-malate, succinate, and acetate (**Figure 2A**)--was not observed in the PaCco and PaCio mutant backgrounds, indicating that the full complement of terminal oxidases is required for this effect. Interestingly, deletion of all non-Cco terminal oxidases revealed just one condition--growth on L-glutamine--in which phenazine production enhanced dye reduction. Characterization of the PaCco and PaCio mutants also revealed several new carbon sources for which phenazines inhibited dye reduction. Together, these results suggest that phenazine production affects tetrazolium dye reduction, and by extension, electron flow through the ETC, differently depending on which terminal oxidases are present.

### Phenazine production affects terminal oxidase gene expression

Previous studies have shown that phenazines oxidize the cellular redox state (5, 23) and that this property affects the activities of regulators that control terminal oxidase expression (11, 13, 24–26). To further characterize the effects of phenazines on the ETC, we created reporter strains corresponding to each of the loci encoding the five major *P. aeruginosa* terminal oxidases (**Figure 3A**). We measured relative expression levels in liquid cultures and biofilms grown on the carbon sources succinate, αKG, glucose, and tryptone. Succinate and αKG constitute two of the four carbon sources that showed phenazine-dependent inhibition of tetrazolium dye reduction in the presence of all terminal oxidases (**Figure 2B**). Glucose and succinate are both popular carbon sources for laboratory media and represent *P. aeruginosa*’s two main pathways for carbon source oxidation, the Entner-Doudoroff pathway and the TCA cycle, respectively (**Figures 1** and **2B**). Tryptone, which is an undefined protein digest, is also a standard carbon source used to grow *P. aeruginosa* in the laboratory and is of interest because it represents a complex condition that may be relevant for *P. aeruginosa* growth in natural settings (27). We found that WT and Δ*phz* showed similar growth kinetics and yields on all of these carbon sources (**Figure S3**). However, we observed slower and more variable growth on glucose than on tryptone, succinate, and αKG, reflecting *P. aeruginosa*’s preference for TCA cycle intermediates over sugars (28). Δ*phz* strains also generally showed a longer lag phase during growth on glucose.

Expression of the *P. aeruginosa* terminal oxidases Cox (or *aa*_3_), Cio, Cco1, and Cco2 during growth in liquid cultures and biofilms is shown in **Figures 4** and **5**, respectively. (We detected expression of *P. aeruginosa*’s fifth major terminal oxidase, Cyo, only in late stationary phase during planktonic growth on tryptone, and at low or negligible levels under all other conditions examined.) For reporter strains grown in liquid cultures, we identified four combinations of reporters and conditions that showed higher expression in the Δ*phz* than in the phenazine-producing background: the Cio and Cco1 reporter strains grown on tryptone, and the Cox and Cio reporter strains grown on αKG. We have previously reported that respiratory chain components involved in nitrate respiration are induced by defects in phenazine production (29), and our observation that the Cox, Cio, and Cco1 terminal oxidases are induced in the Δ*phz* background under some conditions is consistent with this theme. Enhancement of terminal oxidase expression by phenazine deficiency is supported by the notion that they are regulated in response to cellular redox conditions and that phenazines constitute an alternate method of redox balancing when aerobic respiratory activity is limited (5, 30, 31). The higher level of expression for Cox in the phenazine-producing background during growth on tryptone stands out as a unique case and may arise from its control by two independent regulators (i.e., the stationary-phase sigma factor RpoS and the two component repressor system RoxSR) (13, 25).

**Figure 4.**
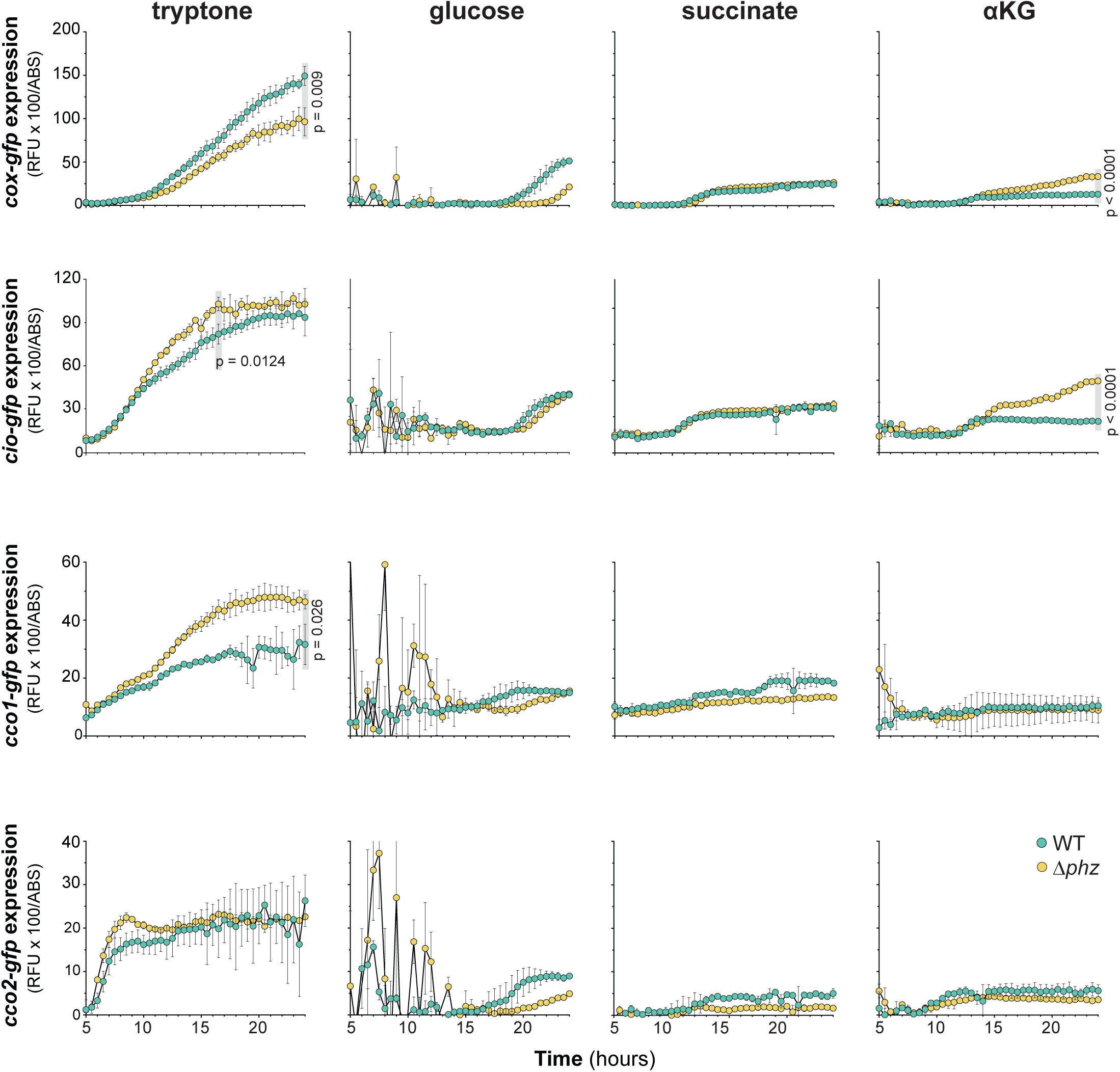
Electron donor and phenazines affect terminal oxidase expression in liquid culture. Mean fluorescence of reporter strains engineered to express GFP under the control of the *cox, cio, cco1*, or *cco2* promoter during liquid culture growth on the indicated carbon sources. Mean fluorescence values were corrected for cell density (OD at 500 nm), and the fluorescence values of a strain expressing GFP without a promoter (the MCS control) were subtracted from each data point. Gray boxes indicate the time points for which p values were determined using unpaired, two-tailed t-tests. Data represent the mean of three biological replicates; error bars denote standard deviation and are not drawn in instances where they would be obscured by point markers.

**Figure 5.**
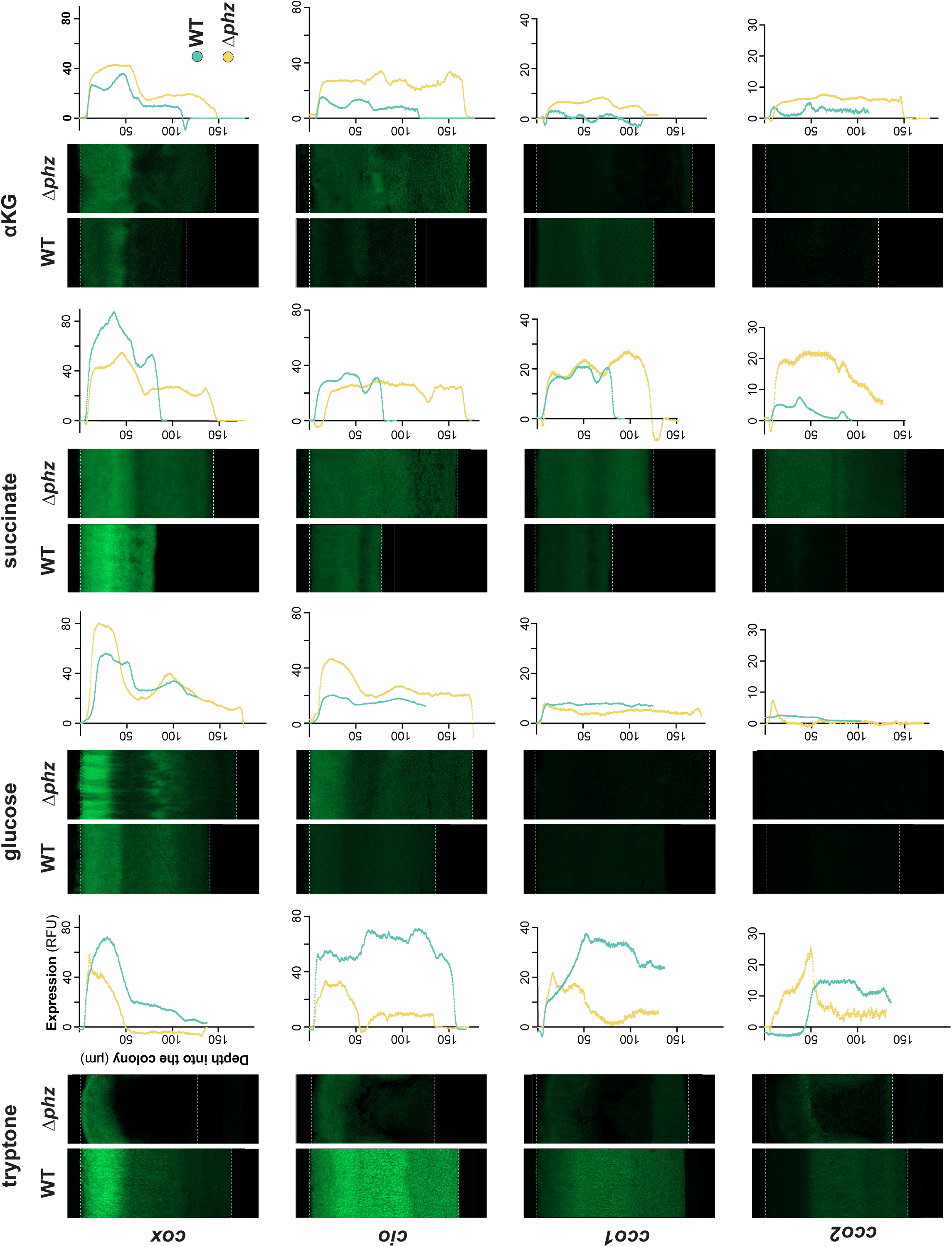
Electron donor and phenazines affect terminal oxidase expression in colony biofilms. Representative images showing expression of GFP under the control of the *cox, cio, cco1*, or *cco2* promoter in thin sections prepared from biofilms grown for three days on the indicated carbon sources Reporter fluorescence is shown in green and is overlain on the DIC image of each colony biofilm. Dotted lines indicate the air-biofilm (top) or agar-biofilm (bottom) interfaces. Graphs show fluorescence values relative to biofilm depth of the corresponding images. Fluorescence values of the MCS control (a strain expressing GFP without a promoter) in either the WT or Δ*phz* background have been subtracted from each plot. The y-axis for each graph provides scale for the respective image, which is representative of at least six biological replicates.

Biofilm growth promotes the formation of resource gradients and spatial heterogeneity that is not observed in liquid culture. We have described two strategies that *P. aeruginosa* can exploit to cope with O_2_ limitation in tryptone-grown biofilms: (1) use of phenazines, which can act as electron shuttles to O_2_ at a distance, and (2) enhanced matrix production that leads to the formation of vertical structures, called “wrinkles”, which increase the overall surface area and therefore cellular access to O_2_ (5, 32, 33). We have previously shown that WT growth on tryptone yields biofilms that are thicker than those formed by Δ*phz*, and attributed the thickness of WT biofilms to the physiological benefit of phenazine production. In this study, however, we found that WT biofilms grown on a defined medium containing glucose, succinate, or αKG were thinner than Δ*phz* biofilms grown under these conditions and that biofilm thickness showed overall variation between conditions. The metabolic pathways that function during growth on distinct carbon sources may differentially affect redox homeostasis and the response to O_2_ limitation. Glucose, for example, can be oxidized through respiration but can also be fermented to lactate, whereas succinate and αKG are not fermentable carbon sources. Finally, we note that the phenazine pyocyanin has been shown to have both beneficial and toxic effects that can influence biofilm development in a condition-dependent manner (34).

Our characterization of terminal oxidase gene expression in biofilms showed several loci and conditions for which expression was enhanced in the absence of phenazines (**Figure 5**), again consistent with complementary roles of the terminal oxidases and phenazines in redox balancing. However, this effect was limited to biofilms grown on defined media containing distinct carbon sources. By contrast, biofilms grown on tryptone consistently showed higher levels of terminal oxidase gene expression in the phenazine-producing background relative to the phenazine-null background. This may be because the greater surface area of tryptone-grown Δ*phz* biofilms increases access to O_2_ and therefore alleviates the need for enhanced terminal oxidase gene expression.

### Phenazine derivatization is highly dependent on mode of growth and carbon source identity

The data presented thus far suggest that PA14’s branched ETC is modulated in response to electron donor identity and the production of phenazines. PA14 makes at least four phenazines (**Figure 6A**), each having a unique redox potential (35, 36). We therefore next asked if the composition of the phenazine pool changes depending on the carbon source. We grew PA14 in liquid cultures and as biofilms on tryptone, glucose, succinate, and αKG and quantified phenazine production using high performance liquid chromatography (HPLC).

**Figure 6.**
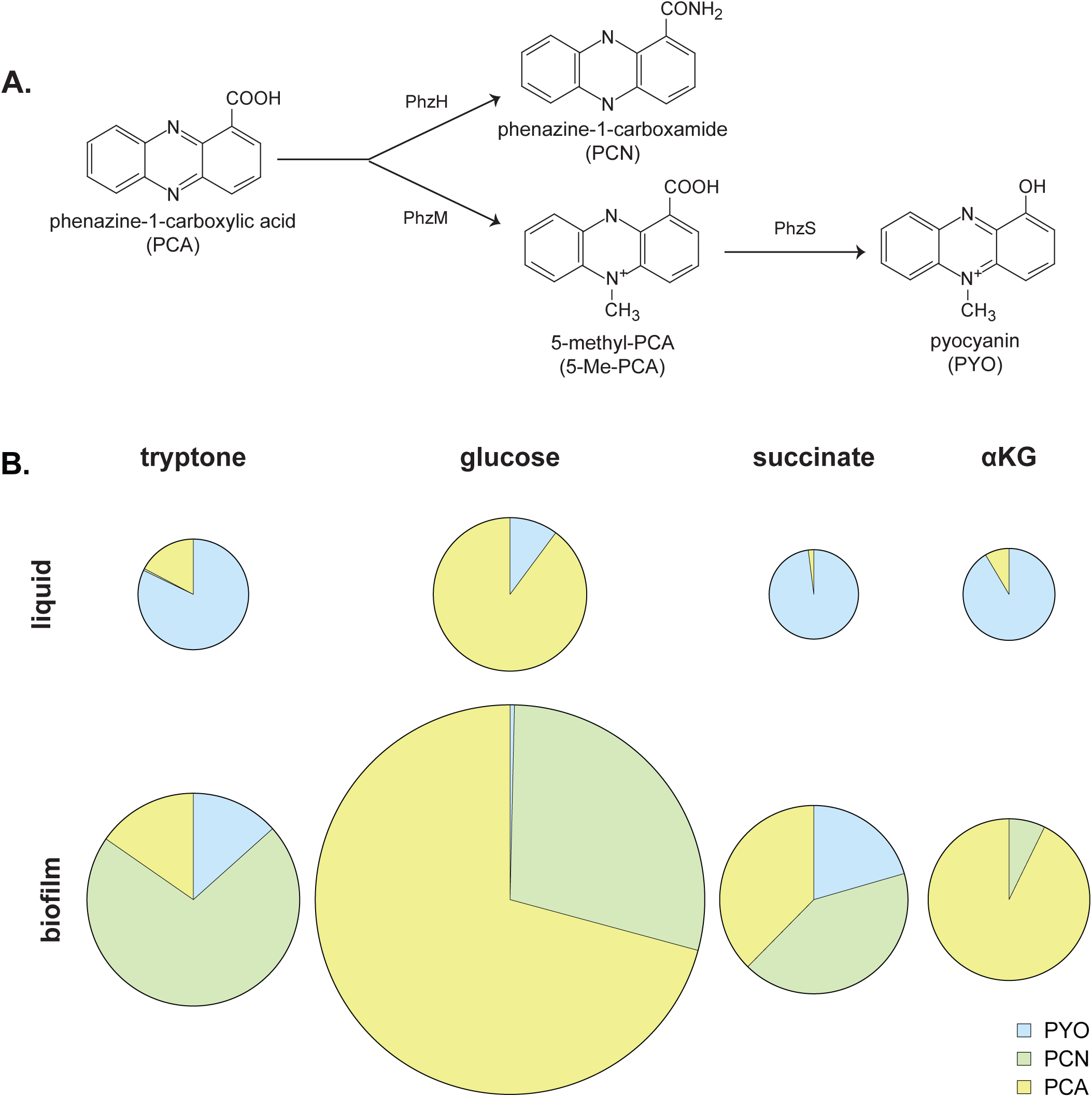
Phenazine production is affected by mode of growth. **(A)** The precursor phenazine phenazine-1-carboxylic acid (PCA) can be modified by PhzH to produce phenazine-1-carboxamide (PCN) and/or PhzM to produce 5-methyl-PCA (5-MCA). 5-MCA can be further modified by PhzS to yield pyocyanin (PYO). **(B)** Phenazines produced by WT PA14 in liquid cultures (top) and biofilms (bottom) grown on the indicated carbon sources. The area of each pie represents the combined concentration of all measured phenazines and each “slice” represents the mean concentration in µM of PCA (yellow), PCN (green), or PYO (blue). Mean values are representative of data from at least six biological replicates. For numerical data, refer to **Figure S4**.

The precursor phenazine, phenazine-1-carboxylic acid (PCA), can be derivatized to phenazine-1-carboxamide (PCN), 5-methyl-PCA (5-Me-PCA), or pyocyanin (PYO) (**Figure 6A**). We focused our analysis on the production of PCA, PCN, and PYO because 5-Me-PCA is highly reactive and unstable (37) and therefore difficult to detect by HPLC. We found a high degree of variation in phenazine production both between carbon sources and liquid culture versus biofilm growth conditions. Most notably, while we found that biofilm growth favored the production of PCN, this phenazine was not detectable in samples from liquid cultures (**Figure 6B**, bottom panel versus top panel, and **Figure S4**). In contrast, while PYO generally contributed the most to the phenazine complement in liquid culture, it was the least-produced phenazine in biofilms. This indicates a general biofilm-specific, but carbon source-independent, switch from PYO to PCN production between the planktonic and biofilm growth states, respectively. However, the extent of PCA derivatization, primarily to PYO (liquid culture) or PCN (biofilm), was dependent on the type of carbon source (**Figure 6B**). These dynamics in phenazine production indicate that the carbon source influences the kinds and amount of phenazines produced by PA14 and highlights the plasticity of phenazine derivatization in response to environmental conditions.

### Biofilms grown on glucose or alpha-ketoglutarate exhibit Cco-independent phenazine reduction

Our observations regarding the condition-dependent effects of phenazines on respiratory activity and terminal oxidase gene expression (**Figures 2-5**), and the high condition-dependence of phenazine derivatization (**Figure 6**), raise the question of how carbon source identity influences phenazine utilization. We have previously shown that phenazine reduction in tryptone-grown biofilms is detectable as a gradient of an increasingly reduced extracellular redox state approaching the biofilm base, and that it requires a Cco terminal oxidase (7). We used redox microelectrodes to measure phenazine reduction in biofilms grown on defined media containing our carbon sources of interest and found that it was detectable during growth on glucose and αKG, but not during growth on succinate (**Figure 7B-D**). Biofilms grown on either glucose and αKG both showed redox profiles that differed from those grown on tryptone: on glucose, phenazines were reduced in the topmost, aerobic zone of the biofilm (**Figure 7B**); on αKG, phenazine reduction was gradual across colony depth and a smaller range of potentials than on either tryptone or glucose (**Figure 7D**). Intriguingly, the Cco oxidases were not required for endogenous phenazine reduction on glucose or αKG. Together, these results indicate that metabolic routes of electron flow to phenazines are altered significantly in response to different carbon sources.

**Figure 7.**
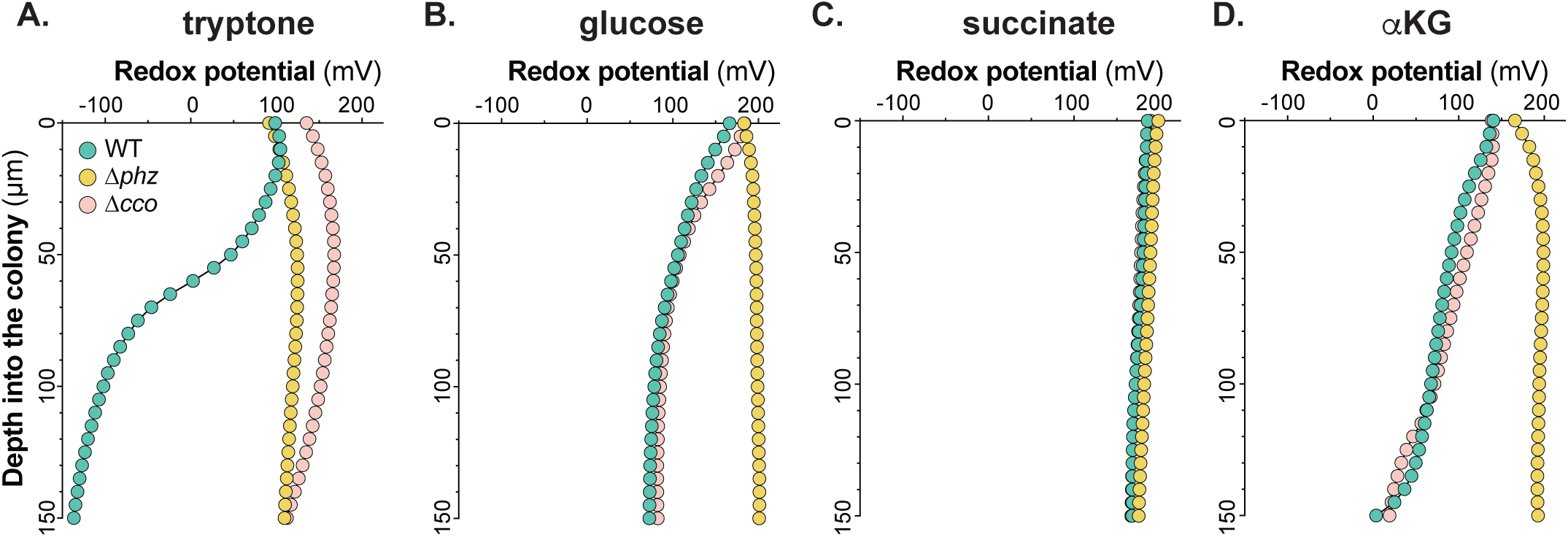
Carbon source alters reduction of endogenously produced phenazines. Change in redox potential across depth for WT, Δ*phz*, and Δ*cco1cco2* (Δ*cco*) colony biofilms grown for three days on tryptone **(A)**, glucose **(B)**, succinate **(C)**, and αKG **(D)**. Data are representative of at least six biological replicates.

### Exogenously provided and endogenous phenazines are subject to distinct reduction mechanisms in biofilms

Because PA14 synthesizes multiple phenazines that have distinct chemical properties (e.g. redox potentials) and may be reduced by distinct mechanisms, we created a “clean” strain background that would allow us to measure the reduction profiles of each phenazine in isolation. To this end, we generated a mutant, referred to as “Δ*HMS*Δ*phz*”, that is unable to produce or modify any of PA14’s known phenazines (**Figure 6A**). We also created the Δ*HMS*Δ*phz*Δ*cco* mutant, which we expected to show defects in phenazine reduction under certain conditions. We then grew these mutants on agar plates each containing one pure phenazine compound and measured phenazine reduction on our carbon sources of interest. Due to the aforementioned instability of 5-Me-PCA, we used its synthetic analog phenazine methosulfate (PMS) as a proxy to study 5-Me-PCA reduction (38). We note that biofilm redox profiles do not show when the microelectrode transitions from the biofilm to the underlying growth medium. We can infer biofilm thicknesses from our characterizations of thin sections (**Figure 5)**, and when comparing the redox profiles obtained for Δ*HMS*Δ*phz* biofilms grown on different conditions, we focused on features most likely to have been present in the biofilms rather than the growth media.

When Δ*HMS*Δ*phz* biofilms were grown on either tryptone or glucose, we found that phenazine reduction activity depended on the methylation state of the derivative, with tryptone-grown biofilms specifically reducing the methylated phenazines PMS and PYO, and glucose-grown biofilms only showing substantial reduction of the non-methylated PCA and PCN (**Figure 8A** and **B**, green traces). While all of these activities were affected, to varying extents, by removal of the Cco terminal oxidases (**Figure 8A and B**, pink traces) the most dramatic effect of the *cco* deletion was abolishment of PMS reduction on tryptone.

**Figure 8.**
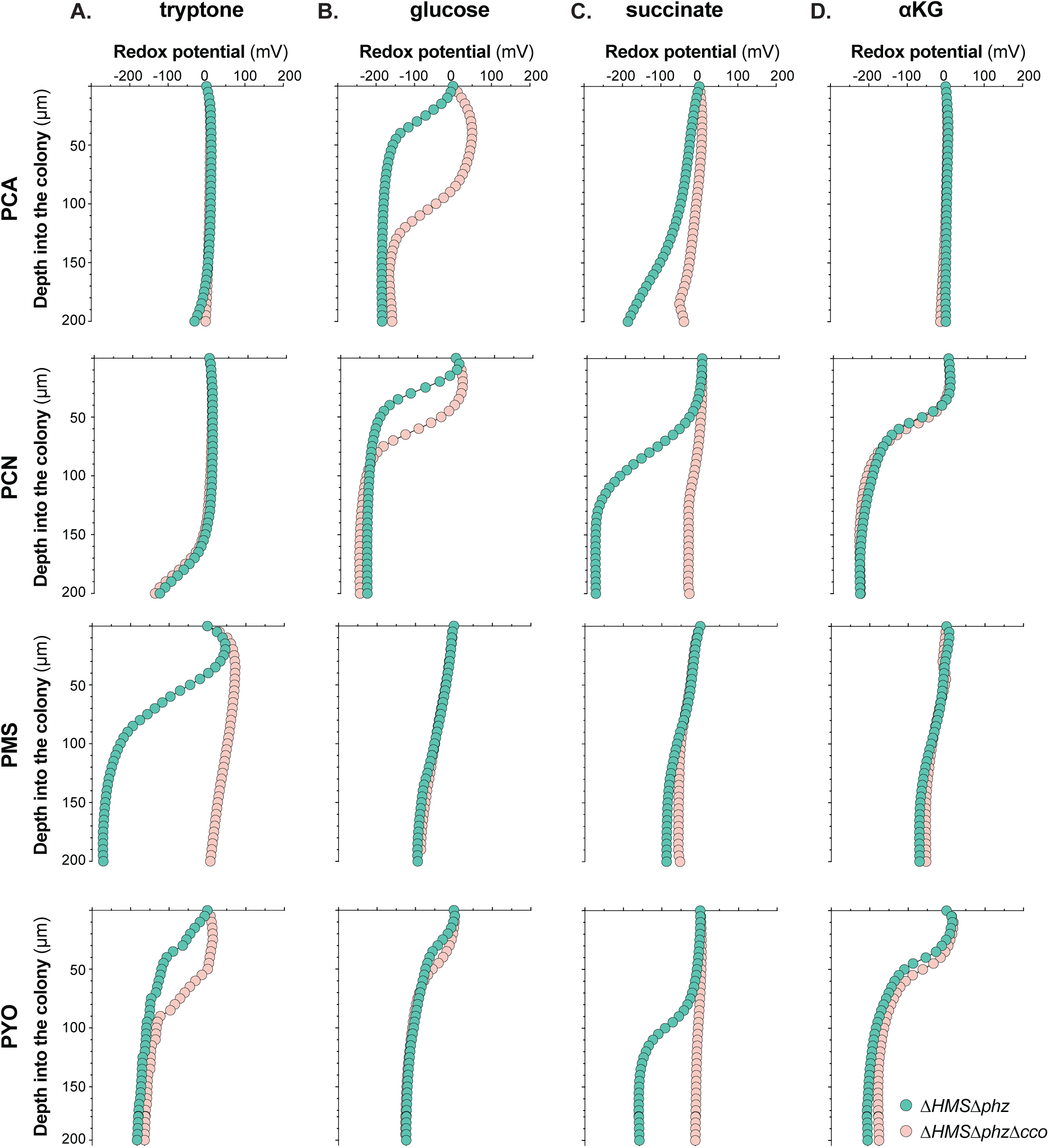
Exogenously provided phenazines exhibit different reduction patterns than those produced endogenously. Change in redox potential across depth for Δ*HMS*Δ*phz* and Δ*HMS*Δ*phz*Δ*cco* colony biofilms grown for three days on tryptone **(A)**, glucose **(B)**, succinate **(C)**, and αKG **(D)** amended with purified phenazine-1-carboxylic acid (PCA), phenazine-1-carboxamide (PCN), phenazine methosulfate (PMS; a synthetic analog of the endogenous phenazine 5-MCA), or pyocyanin (PYO). Data are representative of at least six biological replicates.

Surprisingly, redox profiling of biofilms grown on succinate and added phenazines revealed substantial reduction of PCN and PYO (**Figure 8C**), even though we did not observe reduction of endogenously produced phenazines on this carbon source (**Figure 7C**). This discrepancy could be attributed to (1) an effect of phenazine biosynthesis itself on preferred pathways of carbon source utilization and/or cellular redox state, or (2) different mechanisms of phenazine reduction that operate more readily when phenazines are produced in the cytoplasm. Like that obtained for PMS reduction during growth on tryptone, the phenazine reduction profiles for growth on succinate were also strongly affected by removal of the Cco oxidases. Finally, exogenously provided PCN and PYO were also reduced by biofilm cells growing on αKG (**Figure 8D**). Reduction profiles obtained on this carbon source stand out because they did not depend on the presence of the Cco oxidases, indicating alternate mechanisms of phenazine reduction during growth under this condition. This finding is consistent with the relatively low levels of *cco1* and *cco2* expression we observed for liquid cultures and biofilms grown on αKG (**Figures 4** and **5**), and suggests that phenazine reduction could be mediated by other respiratory chain components or perhaps (an) ETC-independent mechanism(s) on this carbon source (31).

## Concluding remarks

The opportunistic pathogen *P. aeruginosa* is prevalent in healthcare settings, immunocompromised individuals, and people with the inherited disease cystic fibrosis (CF). *P. aeruginosa* is the leading cause of chronic pulmonary infections and morbidity and mortality in patients with CF (3, 39). It exhibits metabolic versatility in its use of diverse carbon sources (i.e., electron donors), its branched aerobic respiratory chain, its capacity for nitrate respiration, and its ability to ferment arginine and pyruvate (9). Studies examining *P. aeruginosa* evolution during CF lung infection have identified mutations associated with changes in nutrient uptake and metabolism (40, 41). Furthermore, ETC genes, including those encoding dehydrogenases and the Cco terminal oxidases, are upregulated in conditions that mimic those found in the CF lung (42). Though *P. aeruginosa*’s metabolic capacities are thought to contribute to its success as a pathogen, an understanding of the condition-dependence and integration of its primary redox pathways is key to interpreting its behavior in hosts.

Beyond canonical metabolic pathways, a further distinctive feature of *P. aeruginosa* is its production of phenazines, small molecules that can carry out extracellular electron shuttling (EET). This property further enhances *P. aeruginosa*’s metabolic versatility because it provides a means to balance the intracellular redox state and generate some ATP when resources such as O_2_ and nitrate are not accessible. *P. aeruginosa* in O_2_-limited environments, such as hypoxic biofilm subzones, can reduce phenazines that then diffuse and transfer electrons to O_2_ or other oxidants available at a distance. Mechanisms supporting EET have been described for diverse bacterial species, including *Shewanella oneidensis, Listeria monocytogenes*, and *Enterococcus faecalis*, and it has been suggested that many more organisms have the potential to carry out this type of metabolism (43–46). EET therefore constitutes a mechanism whereby excreted metabolites can “fine-tune” primary pathways of electron flow and energy generation and thereby promote survival.

Several of our previous studies have highlighted the interdependence of primary metabolism and phenazine-associated physiology (5, 7, 29). In this work, we assessed the effects of phenazine production on respiratory activity and terminal oxidase expression, the effect of growth mode on phenazine production, and the roles of the Cco terminal oxidases in endogenous and exogenous phenazine utilization. We found that the “realms” of *P. aeruginosa* physiology concerning O_2_ and phenazine metabolism strongly influence each other and these effects can differ dramatically depending on carbon source (i.e., electron donor) identity and whether the bacterium is growing in liquid cultures or biofilms. These findings underscore the considerable effect of electron donor(s) on downstream redox balancing mechanisms of *P. aeruginosa*. This work adds to our growing understanding of the different mechanisms that govern *P. aeruginosa* metabolism, which can be modulated precisely to allow for maximal growth and survival under a variety of conditions, including those found within a host.

## Materials and methods

### Strains and growth conditions

*Pseudomonas aeruginosa* strain UCBPP-PA14 (47, 48) precultures were routinely grown for 12-16 hours in 2 ml of lysogeny broth (LB; (49)) in 13 x 100 mm borosilicate glass tubes at 37 °C with shaking at 250 rpm. Precultures serving as biological replicates were inoculated from separate clonal source colonies grown on streaked LB + 1.5% agar plates. Liquid subcultures were made from diluting precultures 1:100 (1:50 for PaCio, phzPaCio, Δ*cco*, and Δ*HMS*Δ*phz*Δ*cco*) in 5 ml of LB and growing in 18 x 150 mm borosilicate glass tubes at 37 °C with shaking at 250 rpm until mid-exponential phase (OD at 500 nm ∼ 0.5). Liquid subcultures were used for experiments unless otherwise noted. Strains used in this study are listed in **Table 1**.

**Table 1:**
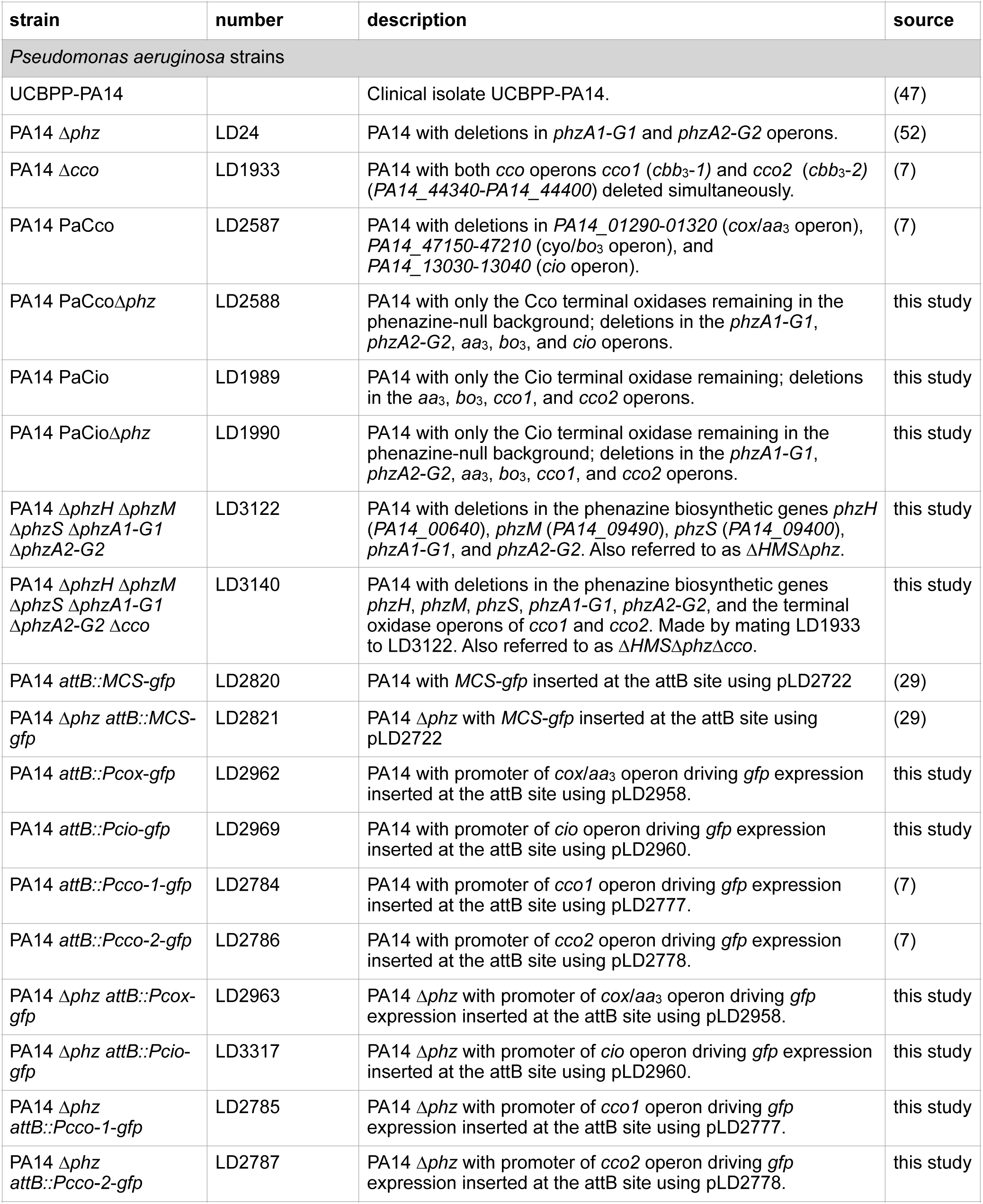
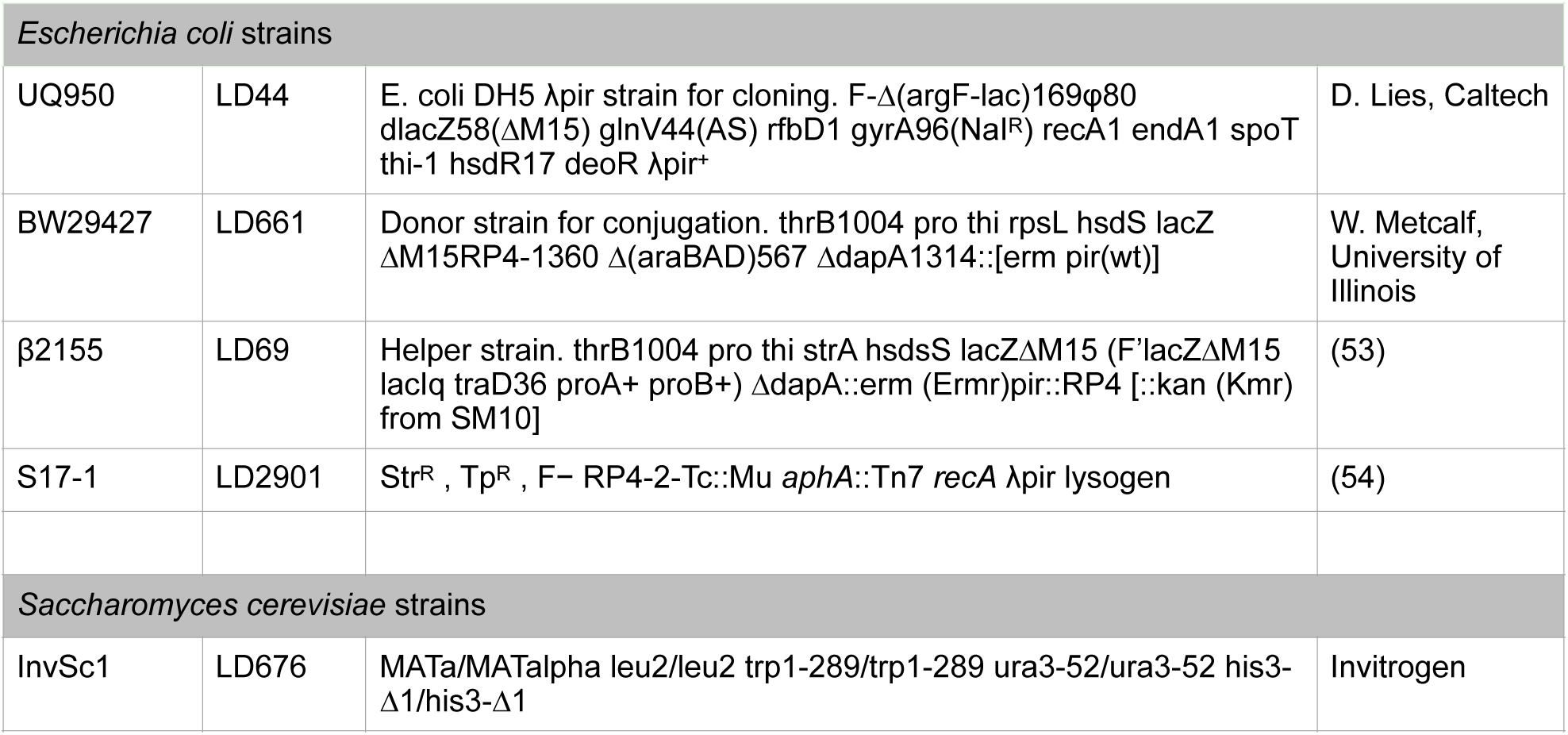
Strains used in this study-1.

The growth media used in this study were 1% tryptone and MOPS minimal medium (50 mM 4-morpholinepropanesulfonic acid (pH 7.2), 43 mM NaCl, 93 mM NH_4_Cl, 2.2 mM KH_2_PO_4_, 1 mM MgSO_4_•7H_2_O, 1 µg/ml FeSO_4_•7H_2_O, and 20 mM carbon source [D-glucose, sodium succinate hexahydrate, or α-ketoglutaric acid sodium salt]). Stock solutions of MgSO_4_, FeSO_4_, and carbon sources were filter sterilized. Agar-containing versions of all growth media contained 1% agar (Teknova A7777).

### *Construction of* P. aeruginosa *deletion mutant strains*

Markerless deletions mutants were created by amplifying ∼ 1 kb of flanking DNA sequence from each side of the gene(s) to be deleted using the primers listed in **Table 2**. These flanking sequences were inserted into pMQ30 via gap repair cloning in *Saccharomyces cerevisiae* strain InvSc1 (50). The resulting plasmids, listed in **Table 3**, were then transformed into *Escherichia coli* strain UQ950, verified by restriction digests and/or sequencing, and moved into PA14 using biparental conjugation. Single recombinants in PA14 were selected using LB agar plates containing 100 µg/ml gentamicin. Markerless deletions (double recombinants) were selected on LB without NaCl containing 10% sucrose and confirmed by PCR. Combinatorial mutants were constructed by using single mutants as hosts for biparental conjugation as described in **Table 1**.

**Table 2:**
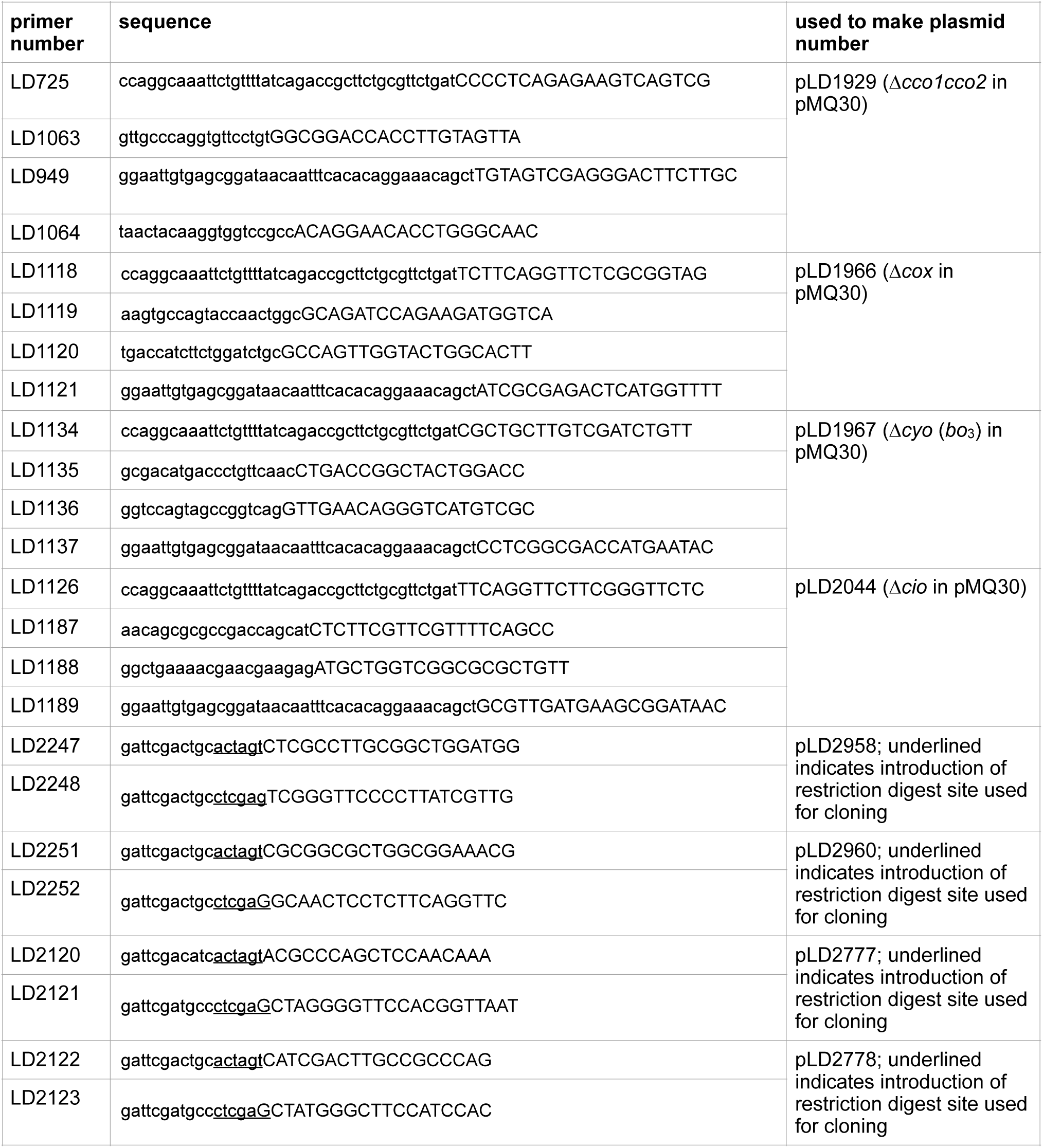
Primers used in this study.

**Table 3:** Plasmids used in this study.

| plasmid | description | source |
| --- | --- | --- |
| pMQ30 | 7.5 kb mobilizable vector; oriT, sacB, Gm<sup>R</sup>. | (50) |
| pLD2722 | Gm<sup>R</sup>, Tet<sup>R</sup> flanked by Flp recombinase target (FRT) sites to resolve out resistance cassettes. | (7) |
| pFLP2 | Flp recombinase-producing plasmid | (55) |
| pLD1929 | $\Delta cco1cco2$ PCR fragment introduced into pMQ30 by gap repair cloning in yeast strain InvSc1. | (7) |
| pLD1966 | $\Delta cox$ (*aa<sub>3</sub>*) PCR fragment introduced into pMQ30 by gap repair cloning in yeast strain InvSc1. | this study |
| pLD1967 | $\Delta cyo$ (*bo<sub>3</sub>*) PCR fragment introduced into pMQ30 by gap repair cloning in yeast strain InvSc1. | this study |
| pLD2044 | $\Delta cio$ PCR fragment introduced into pMQ30 by gap repair cloning in yeast strain InvSc1. | this study |
| pLD2958 | PCR-amplified *cox* (*aa<sub>3</sub>*) promoter ligated into pLD2722 using SpeI and XhoI. | this study |
| pLD2960 | PCR-amplified *cio* promoter ligated into pLD2722 using SpeI and XhoI. | this study |
| pLD2777 | PCR-amplified *cco1* promoter ligated into pLD2722 using SpeI and XhoI. | (7) |
| pLD2778 | PCR-amplified *cco2* promoter ligated into pLD2722 using SpeI and XhoI. | (7) |

### Construction of reporter strains

Reporter constructs for *P. aeruginosa*’s terminal oxidases (Cox, Cio, Cco1, and Cco2) were constructed using the primers listed in **Table 2** to amplify respective promoter regions (500 bp upstream of the respective terminal oxidase operon), adding an SpeI and an XhoI digest site to the 5’ and 3’ ends of the promoter, respectively. Purified PCR products were digested and ligated into the pLD2722 vector at the multiple cloning site (MCS), upstream of the *gfp* sequence. The resulting plasmids (listed in **Table 3**) were transformed into *E. coli* strain UQ950, verified by sequencing, moved into PA14 WT or Δ*phz* using biparental conjugation with *E. coli* strain S17-1. Recombinants were selected for as described previously (7).

### Growth assays and determination of carbon sources that support PA14 growth

One ml of liquid subculture was washed two times in phosphate-buffered saline (PBS; Cold Spring Harbor Laboratories), resuspended in 1 ml MOPS minimal medium without a carbon source, and diluted 1:100 in MOPS minimal medium without a carbon source. One hundred microliters of this cell suspension were dispensed into each well of a phenotype microarray carbon microplate PM1 (Biolog 12111) and incubated at 37 °C with continuous shaking on the medium setting in a Synergy 4 plate reader (BioTek). Growth was measured by taking OD readings at 500 nm every 30 min for 20-24 hr. Each strain was assayed in biological triplicate.

Carbon sources that support growth of PA14 were determined by monitoring growth of WT over 22.5 hours for each biological replicate. After subtracting the cell density (OD at 500 nm) values for the negative control (well A1 on Biolog plate PM1, in which there is no carbon source added), a cutoff of 0.15 was applied to the other 95 cell density values; anything falling below this cutoff was designated as background. In the rare instances where there was a discrepancy between the biological replicates, the growth curves of each replicate was manually inspected to determine whether there was growth or not. We found that 32 of 95 carbon sources in Biolog plate PM1 allowed for growth of PA14 WT (**Supplementary File 1**). We ensured that only those carbon sources that support growth of WT also allow the terminal oxidase and/or phenazine mutants to grow by visually inspecting their growth.

### Determination of carbon sources that support reduction of tetrazolium violet

For dye reduction assays, PA14 cells were prepared and assayed according to manufacturer’s instructions. Briefly, PBS-washed cells were resuspended in 1x IF-0 (IF-0a GN/GP Base Inoculating Fluid, Biolog 72268) to a percent transmittance of 42. The 42 %T cell suspension was diluted 1:5 in 1x IF-0 + 1x Redox Dye A (tetrazolium violet; Biolog 74221). One hundred microliters of this cell + dye suspension were dispensed into each well of a PM1 microplate and incubated at 37 °C with continuous shaking on the medium setting in a Synergy 4 plate reader (BioTek). Dye reduction was measured by taking absorbance readings at 590 nm, as per manufacturer’s instructions, every 30 min for 20-24 hr. Absorbance values for a well containing no carbon source (A1) were subtracted from each data point. Tetrazolium dye reduction was only determined for carbon sources that supported PA14 growth (**Supplementary File S1** and **Figure S2**). Dye reduction patterns of WT, PaCco, and PaCio were compared to their respective phenazine-null counterparts and the effect of phenazines on dye reduction in each carbon source was determined (**Figure 3B**).

### Liquid culture growth and terminal oxidase reporter expression assays

One ml of liquid subcultures was washed two times in PBS and resuspended in 1 ml PBS. These cells were then diluted 1:100 in respective growth medium in a clear, flat-bottom polystyrene black 96-well plate (VWR 82050-756) and incubated at 37 °C with continuous shaking on the medium setting in a Synergy 4 plate reader (BioTek). Growth was monitored by taking OD readings at 500 nm and *gfp* expression was assessed by taking fluorescence readings at excitation and emission wavelengths of 480 nm and 510 nm, respectively, every 30 minutes for 20-24 hr. Fluorescence and absorbance readings were taken simultaneously. For growth curves (**Figure S4**), OD at 500 nm values of a growth medium blank was subtracted from the growth values of each strain. For expression data shown in **Figure 4**, RFU values of each reporter were adjusted for cell density by dividing the fluorescence value at each time point with the corresponding cell density values. RFU values, also corrected for cell density, for a strain lacking a promoter upstream of the *gfp* gene (MCS-*gfp*) were subtracted from the fluorescence values of each reporter.

### Thin sectioning assays

Thin sectioning assays were performed as described previously (Cornell et al. 2018). Briefly, 5 µl of subcultures were spotted onto 1% agar plates containing two-layered growth medium (1% tryptone, MOPS-glucose, MOPS-succinate, or MOPS-MOPS-αKG) and grown in the dark at 25 °C with > 90% humidity (Percival CU-22L). After three days, colonies were covered with an agar layer and sandwiched colonies were lifted from the bottom layer, washed for 10 min in PBS (pH 7.4) at room temperature in the dark, and fixed in 4% paraformaldehyde in PBS overnight at room temperature in the dark. Fixed colonies were washed twice in PBS and dehydrated through a series of ethanol washes. Colonies were cleared via three 60-min incubations in Histoclear-II (National Diagnostics HS-202) and infiltrated with wax via two separate washes of 100% Paraplast Xtra paraffin wax (Thermo Fisher Scientific 50-276-89) for 2 hr each at 55 °C, then colonies were allowed to polymerize overnight at 4 °C. Tissue processing was performed using an STP120 Tissue Processor (Thermo Fisher Scientific 813150). Trimmed blocks were sectioned in 10-µm-thick sections perpendicular to the plane of the colony using an automatic microtome (Thermo Fisher Scientific 905200ER) and collected onto slides. Slides were air-dried overnight, heat-fixed on a hotplate for 1 hr at 45 °C, and rehydrated. Rehydrated colonies were immediately mounted in TRIS-Buffered DAPI:Fluorogel (Thermo Fisher Scientific 50-246-93) and overlaid with a coverslip. Differential interference contrast (DIC) and fluorescent confocal images were captured from using an LSM800 confocal microscope (Zeiss, Germany). n ≥ 6 for each strain on each carbon source.

### Thin sectioning data analysis

Fluorescence intensities of terminal-oxidase GFP reporters expressed in biofilm thin sections were determined using Fiji image processing software (51) and plotted after subtracting values for the respective strains containing a promoterless GFP reporter. Cropped versions of the analyzed images are shown. Background fluorescence was removed by aligning images for terminal-oxidase GFP reporters with those for the promoterless GFP reporter and subtracting their fluorescence values using Adobe Photoshop CC.

### Phenazine detection and quantification

(i) For phenazine extraction from liquid cultures, 1 ml of liquid subcultures was washed three times in 1x PBS and resuspended in 1 ml 1x PBS. Washed cells were diluted 1:100 in 200 µl of respective growth medium in a clear, flat-bottom polystyrene 96-well plate (VWR 82050-716) and incubated at 37 °C with continuous shaking on the medium setting in a Synergy 4 plate reader (BioTek). Growth was measured by taking OD readings at 500 nm every 30 min and cultures were monitored to determine onset of stationary phase. Five hours after each culture reached stationary phase, samples were collected into a microfuge tube and centrifuged for two minutes at 14,000 rpm to pellet cells. The supernantant of each sample was applied to a 0.22 µm Spin-X column (VWR 29442-754), centrifuged for two minutes at 14,000 rpm, and 200 µl of the resulting cell-free flow-through was loaded into an HPLC vial for analysis. For each growth condition, n = 10.

(ii) For phenazine extraction from biofilms, 1 ml of liquid subcultures was washed twice in 1x PBS and resuspended in 1 ml of 1x PBS. A 25 mm filter disk with a pore size of 0.2 µm (GE Healthcare 110606) was placed into the center of one 35 x 10 mm round Petri dish (VWR 25373-041) filled with 4 ml of 1% tryptone agar, MOPS-glucose agar, MOPS-succinate agar, or MOPS-αKG agar and 10 µl of the washed cells were spotted onto the filter disk. Colony biofilms were grown for 3 days in the dark at 25 °C with > 90% humidity, after which point each colony and filter were lifted off their plate. The agar upon which the biofilm had developed was placed into a 5-ml aliquot of 100% methanol in a polypropylene conical tube, and phenazines were extracted from the agar overnight at room temperature in the dark. Three hundred microliters of the phenazine extraction were filtered through a 0.22 µm Spin-X column as described above and 200 µl of the cell-free flow-through were loaded into an HPLC vial for analysis. For each growth condition, n ≥ 6.

(iii) Phenazines were identified using high-performance liquid chromatography (Agilent 1100 HPLC System) as described previously (38, 52) and comparing sample peaks to peaks of pure phenazine standards run as controls. The height of each peak (AU) was used to determine the concentrations of each phenazine. For phenazine quantification from biofilms, the calculated phenazine concentration was multiplied by 1.25 to account for dilution during the phenazine extraction step. Dilution standards of purified PYO, PCA, and PCN were prepared at known concentrations and extinction coefficients (ε) were generated for each: ε_PYO_ = 1.34 µM/AU, ε_PCA_ = 7.686 µM/AU, ε_PCN_ = 7.518 µM/AU.

### Redox microprofiling

(i) Extracellular redox states of day 3 (∼ 72 hr) phenazine-producing biofilms were measured using a 25 µm-tip redox microelectrode and external reference (Unisense RD25 and REF-RM). Colony biofilms were grown by spotting 10 µl of liquid subcultures onto 60 ml of growth medium (1% tryptone, MOPS-glucose, MOPS-succinate, or MOPS-αKG agar) in a 10 cm × 10 cm × 1.5 cm square Petri dish (LDP D210-16). Calibration and redox measurements were performed using the SensorTrace Profiling software (Unisense) as described previously (7).

(ii) For measuring extracellular redox states of biofilms grown on exogenously added phenazines, purified phenazines compounds were added to the respective growth media (described above) to final concentrations of 200 µM after cooling to ∼ 55 °C in a water bath. The phenazines tested were PCA (Apexmol), PCN (Apexmol), PMS (Thermo Fisher Scientific AC130160010), and PYO (Cayman Chemicals). PCA and PCN stock solutions were 25 mM in DMSO while PYO was 200 mM in DMSO. The PMS stock solution was 200 mM in water. Extracellular redox states of biofilms were measured on DMSO-only plates to ensure that the solvent did not alter extracellular redox state. Extracellular redox states of day 3 (∼ 72 hr) biofilms grown on exogenously added phenazines were measured as above. Redox potential values at the surface of the biofilm (0 µm) were set to 0 mV and remaining values for each plot were adjusted accordingly to facilitate comparisons of phenazine reduction patterns between conditions.

## Acknowledgements

This work was supported by NIH/NIAID grant R01AI103369, an NSF Career Award to L.E.P.D., and NIH training grant 5T32GM008798 to J.J.

## Legends

**Figure S1. Growth of WT and Δ*phz* in phenotype microarray plates**. 32 of the provided 95 carbon sources supported growth of PA14 strains containing the full complement of terminal oxidases (WT and Δ*phz*). Each trace corresponds to a biological replicate, with WT shown in green and Δ*phz* shown in yellow. Growth was assessed by measuring OD at 500 nm every 30 minutes for 20-24 hours.

**Figure S2. Phenazines influence tetrazolium dye reduction**. Dye reduction **(A)** and growth **(B)** patterns of WT (green) and Δ*phz* (yellow) strains in succinate, D,L-malate, acetate, and α-ketoglutarate (αKG). Each trace corresponds to a biological replicate, and representative curves are shown in **Figure 2A**.

**Figure S3. Growth curves of terminal oxidase reporters**. Mean cell density values of reporter strains engineered to express GFP under the control of the *cox, cio, cco1*, or *cco2* terminal oxidase promoter during liquid culture growth on 1% tryptone, glucose, succinate, or αKG. Corresponding reporter expression data are shown in **Figure 4**. Data represent the mean of three biological replicates and error bars denote standard deviation and are not drawn in instances where they would be obscured by point markers.

**Figure S4. Phenazine production by *P. aeruginosa* in carbon sources of interest**. Concentrations (µM) of phenazine-1-carboxylic acid (PCA), phenazine-1-carboxamide, green (PCN), and pyocyanin (PYO) produced by WT on 1% tryptone, MOPS-glucose, MOPS-succinate, and MOPS-αKG. Mean values are represented in **Figure 6**.

**Supplementary File S1. *P. aeruginosa* growth in phenotype microarray plates**. Raw data for growth of WT, Δ*phz*, PaCco, PaCcoΔ*phz*, PaCio, PaCioΔ*phz* in Biolog phenotype microarray plate PM1. Cells were grown in MOPS-buffered medium with a different, manufacturer-provided carbon source in each well. Well A1 contains no carbon source and serves as the negative control. OD values at 500 nm were taken every 30 minutes for 20-24 hours. Each strain was assayed in biological triplicate.

**Supplementary File S2. *P. aeruginosa* dye reduction in phenotype microarray plates**. Raw data for tetrazolium dye reduction of WT, Δ*phz*, PaCco, PaCcoΔ*phz*, PaCio, PaCioΔ*phz* in Biolog phenotype microarray plate PM1. Cells were grown in manufacturer-provided “IF” growth medium supplemented with tetrazolium dye with a different carbon source in each well. Well A1 contains no carbon source and serves as the negative control. Absorbance values at 590 nm were taken every 30 minutes for 20-24 hours. Each strain was assayed in biological triplicate.

## References

1. Madigan MT, Martinko JM, Bender KS, Buckley DH, Stahl DA. 2015. Brock Biology of Microorganisms. Pearson.

2. Poole RK, Cook GM. 2000. Redundancy of aerobic respiratory chains in bacteria? Routes, reasons and regulation. Adv Microb Physiol 43:165–224.

3. Høiby N, Ciofu O, Johansen HK, Song Z-J, Moser C, Jensen PØ, Molin S, Givskov M, Tolker-Nielsen T, Bjarnsholt T. 2011. The clinical impact of bacterial biofilms. Int J Oral Sci 3:55–65.

4. Elias S, Banin E. 2012. Multi-species biofilms: living with friendly neighbors. FEMS Microbiol Rev 36:990–1004.

5. Dietrich LEP, Okegbe C, Price-Whelan A, Sakhtah H, Hunter RC, Newman DK. 2013. Bacterial community morphogenesis is intimately linked to the intracellular redox state. J Bacteriol 195:1371–1380.

6. Kempes CP, Okegbe C, Mears-Clarke Z, Follows MJ, Dietrich LEP. 2014. Morphological optimization for access to dual oxidants in biofilms. Proc Natl Acad Sci U S A 111:208–213.

7. Jo J, Cortez KL, Cornell WC, Price-Whelan A, Dietrich LE. 2017. An orphan cbb3-type cytochrome oxidase subunit supports Pseudomonas aeruginosa biofilm growth and virulence. Elife 6.

8. White D, Drummond JT, Fuqua C. 2012. The Physiology and Biochemistry of Prokaryotes. Oxford University Press.

9. Williams HD, Zlosnik JEA, Ryall B. 2007. Oxygen, cyanide and energy generation in the cystic fibrosis pathogen Pseudomonas aeruginosa. Adv Microb Physiol 52:1–71.

10. Arai H. 2011. Regulation and Function of Versatile Aerobic and Anaerobic Respiratory Metabolism in Pseudomonas aeruginosa. Front Microbiol 2:103.

11. Comolli JC, Donohue TJ. 2004. Differences in two Pseudomonas aeruginosa cbb3 cytochrome oxidases. Mol Microbiol 51:1193–1203.

12. Alvarez-Ortega C, Harwood CS. 2007. Responses of Pseudomonas aeruginosato low oxygen indicate that growth in the cystic fibrosis lung is by aerobic respiration. Mol Microbiol 65:153–165.

13. Kawakami T, Kuroki M, Ishii M, Igarashi Y, Arai H. 2010. Differential expression of multiple terminal oxidases for aerobic respiration in Pseudomonas aeruginosa. Environ Microbiol 12:1399–1412.

14. Arai H, Kawakami T, Osamura T, Hirai T, Sakai Y, Ishii M. 2014. Enzymatic characterization and in vivo function of five terminal oxidases in Pseudomonas aeruginosa. J Bacteriol 196:4206–4215.

15. Hirai T, Osamura T, Ishii M, Arai H. 2016. Expression of multiple cbb3 cytochrome c oxidase isoforms by combinations of multiple isosubunits in Pseudomonas aeruginosa. Proc Natl Acad Sci U S A.

16. Aussel L, Pierrel F, Loiseau L, Lombard M, Fontecave M, Barras F. 2014. Biosynthesis and physiology of coenzyme Q in bacteria. Biochim Biophys Acta 1837:1004–1011.

17. Saint-Amans S, Girbal L, Andrade J, Ahrens K, Soucaille P. 2001. Regulation of carbon and electron flow in Clostridium butyricum VPI 3266 grown on glucose-glycerol mixtures. J Bacteriol 183:1748–1754.

18. Otten MF, Stork DM, Reijnders WN, Westerhoff HV, Van Spanning RJ. 2001. Regulation of expression of terminal oxidases in Paracoccus denitrificans. Eur J Biochem 268:2486–2497.

19. Berridge MV, Herst PM, Tan AS. 2005. Tetrazolium dyes as tools in cell biology: new insights into their cellular reduction. Biotechnol Annu Rev 11:127–152.

20. Tachon S, Michelon D, Chambellon E, Cantonnet M, Mezange C, Henno L, Cachon R, Yvon M. 2009. Experimental conditions affect the site of tetrazolium violet reduction in the electron transport chain of Lactococcus lactis. Microbiology 155:2941–2948.

21. Pierson LS 3rd, Thomashow LS. 1992. Cloning and heterologous expression of the phenazine biosynthetic locus from Pseudomonas aureofaciens 30-84. Mol Plant Microbe Interact 5:330–339.

22. Culbertson JE, Toney MD. 2013. Expression and characterization of PhzE from P. aeruginosa PAO1: aminodeoxyisochorismate synthase involved in pyocyanin and phenazine-1-carboxylate production. Biochim Biophys Acta 1834:240–246.

23. Price-Whelan A, Dietrich LEP, Newman DK. 2007. Pyocyanin alters redox homeostasis and carbon flux through central metabolic pathways in Pseudomonas aeruginosa PA14. J Bacteriol 189:6372–6381.

24. Comolli JC, Donohue TJ. 2002. Pseudomonas aeruginosa RoxR, a response regulator related to Rhodobacter sphaeroides PrrA, activates expression of the cyanide-insensitive terminal oxidase. Mol Microbiol 45:755–768.

25. Osamura T, Kawakami T, Kido R, Ishii M, Arai H. 2017. Specific expression and function of the A-type cytochrome c oxidase under starvation conditions in Pseudomonas aeruginosa. PLoS One 12:e0177957.

26. Lin Y-C, Cornell WC, Jo J, Price-Whelan A, Dietrich LEP. 2018. The Pseudomonas aeruginosa Complement of Lactate Dehydrogenases Enables Use of d- and l-Lactate and Metabolic Cross-Feeding. MBio 9.

27. Frimmersdorf E, Horatzek S, Pelnikevich A, Wiehlmann L, Schomburg D. 2010. How Pseudomonas aeruginosa adapts to various environments: a metabolomic approach. Environ Microbiol 12:1734–1747.

28. Rojo F. 2010. Carbon catabolite repression in Pseudomonas : optimizing metabolic versatility and interactions with the environment. FEMS Microbiol Rev 34:658–684.

29. Lin Y-C, Sekedat MD, Cornell WC, Silva GM, Okegbe C, Price-Whelan A, Vogel C, Dietrich LEP. 2018. Phenazines Regulate Nap-Dependent Denitrification in Pseudomonas aeruginosa Biofilms. J Bacteriol 200.

30. Wang Y, Kern SE, Newman DK. 2010. Endogenous phenazine antibiotics promote anaerobic survival of Pseudomonas aeruginosa via extracellular electron transfer. J Bacteriol 192:365–369.

31. Glasser NR, Kern SE, Newman DK. 2014. Phenazine redox cycling enhances anaerobic survival in Pseudomonas aeruginosa by facilitating generation of ATP and a proton-motive force. Mol Microbiol 92:399–412.

32. Okegbe C, Price-Whelan A, Dietrich LEP. 2014. Redox-driven regulation of microbial community morphogenesis. Curr Opin Microbiol 18:39–45.

33. Okegbe C, Fields BL, Cole SJ, Beierschmitt C, Morgan CJ, Price-Whelan A, Stewart RC, Lee VT, Dietrich LEP. 2017. Electron-shuttling antibiotics structure bacterial communities by modulating cellular levels of c-di-GMP. Proc Natl Acad Sci U S A 114:E5236–E5245.

34. Meirelles LA, Newman DK. 2018. Both toxic and beneficial effects of pyocyanin contribute to the lifecycle of Pseudomonas aeruginosa. Mol Microbiol.

35. Price-Whelan A, Dietrich LEP, Newman DK. 2006. Rethinking “secondary” metabolism: physiological roles for phenazine antibiotics. Nat Chem Biol 2:71–78.

36. Wang Y, Newman DK. 2008. Redox reactions of phenazine antibiotics with ferric (hydr)oxides and molecular oxygen. Environ Sci Technol 42:2380–2386.

37. Hansford GS, Holliman FG, Herbert RB. 1972. Pigments of Pseudomonas species. IV In vitro and in vivo conversion of 5-methylphenazinium-1-carboxylate into aeruginosin A. J Chem Soc Perkin 1 1:103–105.

38. Sakhtah H, Koyama L, Zhang Y, Morales DK, Fields BL, Price-Whelan A, Hogan DA, Shepard K, Dietrich LEP. 2016. The Pseudomonas aeruginosa efflux pump MexGHI-OpmD transports a natural phenazine that controls gene expression and biofilm development. Proc Natl Acad Sci U S A 113:E3538–47.

39. Williams HD, Davies JC. 2012. Basic science for the chest physician: Pseudomonas aeruginosa and the cystic fibrosis airway. Thorax 67:465–467.

40. Son MS, Matthews WJ Jr, Kang Y, Nguyen DT, Hoang TT. 2007. In vivo evidence of Pseudomonas aeruginosa nutrient acquisition and pathogenesis in the lungs of cystic fibrosis patients. Infect Immun 75:5313–5324.

41. Hoboth C, Hoffmann R, Eichner A, Henke C, Schmoldt S, Imhof A, Heesemann J, Hogardt M. 2009. Dynamics of adaptive microevolution of hypermutable Pseudomonas aeruginosa during chronic pulmonary infection in patients with cystic fibrosis. J Infect Dis 200:118–130.

42. Eichner A, Günther N, Arnold M, Schobert M, Heesemann J, Hogardt M. 2014. Marker genes for the metabolic adaptation of Pseudomonas aeruginosa to the hypoxic cystic fibrosis lung environment. Int J Med Microbiol 304:1050–1061.

43. Kotloski NJ, Gralnick JA. 2013. Flavin electron shuttles dominate extracellular electron transfer by Shewanella oneidensis. MBio 4.

44. Mevers E, Su L, Pishchany G, Baruch M, Cornejo J, Hobert E, Dimise E, Ajo-Franklin CM, Clardy J. 2019. An elusive electron shuttle from a facultative anaerobe. Elife 8.

45. Light SH, Su L, Rivera-Lugo R, Cornejo JA, Louie A, Iavarone AT, Ajo-Franklin CM, Portnoy DA. 2018. A flavin-based extracellular electron transfer mechanism in diverse Gram-positive bacteria. Nature 562:140–144.

46. Keogh D, Lam LN, Doyle LE, Matysik A, Pavagadhi S, Umashankar S, Low PM, Dale JL, Song Y, Ng SP, Boothroyd CB, Dunny GM, Swarup S, Williams RBH, Marsili E, Kline KA. 2018. Extracellular Electron Transfer Powers Enterococcus faecalis Biofilm Metabolism. MBio 9.

47. Rahme LG, Stevens EJ, Wolfort SF, Shao J, Tompkins RG, Ausubel FM. 1995. Common virulence factors for bacterial pathogenicity in plants and animals. Science 268:1899–1902.

48. Mathee K. 2018. Forensic investigation into the origin of Pseudomonas aeruginosa PA14 - old but not lost. J Med Microbiol 67:1019–1021.

49. Bertani G. 2004. Lysogeny at mid-twentieth century: P1, P2, and other experimental systems. J Bacteriol 186:595–600.

50. Shanks RMQ, Caiazza NC, Hinsa SM, Toutain CM, O’Toole GA. 2006. Saccharomyces cerevisiae-based molecular tool kit for manipulation of genes from gram-negative bacteria. Appl Environ Microbiol 72:5027–5036.

51. Schindelin J, Arganda-Carreras I, Frise E, Kaynig V, Longair M, Pietzsch T, Preibisch S, Rueden C, Saalfeld S, Schmid B, Tinevez J-Y, White DJ, Hartenstein V, Eliceiri K, Tomancak P, Cardona A. 2012. Fiji: an open-source platform for biological-image analysis. Nat Methods 9:676–682.

52. Dietrich LEP, Price-Whelan A, Petersen A, Whiteley M, Newman DK. 2006. The phenazine pyocyanin is a terminal signalling factor in the quorum sensing network of Pseudomonas aeruginosa. Mol Microbiol 61:1308–1321.

53. Dehio C, Meyer M. 1997. Maintenance of broad-host-range incompatibility group P and group Q plasmids and transposition of Tn5 in Bartonella henselae following conjugal plasmid transfer from Escherichia coli. J Bacteriol 179:538–540.

54. Teng F, Murray BE, Weinstock GM. 1998. Conjugal transfer of plasmid DNA from Escherichia coli to enterococci: a method to make insertion mutations. Plasmid 39:182–186.

55. Hoang TT, Karkhoff-Schweizer RR, Kutchma AJ, Schweizer HP. 1998. A broad-host-range Flp-FRT recombination system for site-specific excision of chromosomally-located DNA sequences: application for isolation of unmarked Pseudomonas aeruginosa mutants. Gene 212:77–86.

